# Virus-like particle vaccines targeting a key epitope in circumsporozoite protein provide sterilizing immunity against malaria

**DOI:** 10.1101/2025.03.03.641239

**Authors:** Yogesh Nepal, Alexandra Francian, Yevel Flores-Garcia, Bryce T. Roberts, Sunil A. David, Fidel Zavala, Bryce Chackerian

## Abstract

Vaccines that target the pre-erythrocytic stage of the malaria lifecycle have the potential to provide sterilizing immunity but must elicit sustained, high-titer antibody responses to completely prevent infection. Most pre-erythrocytic vaccines target circumsporozoite protein (CSP), the major surface antigen on *Plasmodium falciparum* sporozoites. Antibodies targeting distinct epitopes within the central repeat region of CSP have the potential to provide protection from infection, but we have focused on developing vaccines that target a highly vulnerable CSP epitope that is targeted by the potent monoclonal antibody L9. In a previous study, we produced a pre-erythrocytic vaccine displaying a synthetic peptide representing the L9 epitope on Qβ bacteriophage virus-like particles (VLPs). This vaccine elicited strong anti-CSP antibody responses that protected mice from malaria challenge. Here, we asked whether the structural context of the L9 epitope influences the quality of antibody responses. We compared the immunogenicity and protective efficacy of Qβ L9 VLPs to recombinant VLPs that display the L9 peptide in a structure that is hypothesized to mimic its native conformation. Recombinant MS2 bacteriophage VLPs displaying various lengths of the L9 epitope were produced and immunogenicity and protective efficacy were evaluated in mice. Our results demonstrate that MS2 L9 VLPs, particularly those displaying longer L9 peptides and in combination with a potent novel adjuvant, elicit strong and durable antibody responses that lower malaria liver burden and prevent infection. We also compared the efficacy of L9-targeted vaccines to the licensed vaccine, RTS,S/AS01_E_ (Mosquirix™, GSK). Immunization with Qβ L9 VLPs, MS2 L9 VLPs, and RTS,S/AS01_E_ provided significant protection from liver-stage infection in a mouse model; immunization with Qβ L9 VLPs elicited sterilizing immunity in the highest percentage of mice. A combination vaccine consisting of MS2 L9 and Qβ L9 VLPs, each presenting the L9 epitope in distinct structural forms, provided the strongest protection, reducing liver parasite burden and promoting sterilizing immunity more effectively than the licensed RTS,S/AS01_E_ vaccine.

## INTRODUCTION

*Plasmodium* parasite infections cause more than 200 million cases of malaria each year, resulting in over 600,000 deaths^1^. Malaria cases are continuing to increase due to the expanding range of mosquito vectors, the growing number of insecticide-resistant mosquitoes, and the emergence of drug-resistant *Plasmodium* strains^2–5^. However, the implementation of the first WHO-approved vaccines targeting *Plasmodium falciparum* (*Pf*), the predominant species of *Plasmodium* that causes severe disease in humans, is an opportunity to reverse this trend. RTS,S/AS01_E_ (Mosquirix™; GSK) and R21/Matrix-M are vaccines that target the pre-erythrocytic stage of the *Pf* lifecycle, during which sporozoites injected by infected mosquitoes transit from the skin to the liver to establish infection. Both vaccines target the circumsporozoite protein (CSP), a protein that coats the surface of the infectious sporozoite and is involved in parasite motility and hepatocyte invasion^6–8^. Vaccines targeting CSP are appealing because they can potentially provide sterilizing immunity by preventing the parasite from establishing infection prior to the symptomatic blood stage. However, completely preventing liver infection comes with several challenges. Sporozoites can reach the liver within an hour of transmission by an infected mosquito^9^. Moreover, a single infected hepatocyte is sufficient to establish a full-blown blood-stage infection^10^. Because of this short window of vulnerability and the high bar for providing sterilizing immunity, effective malaria vaccines that target CSP must elicit remarkably high-titer and high-quality antibody responses at levels that are sustained to the greatest extent possible.

These factors may explain the limitations of first-generation malaria vaccines. In order to elicit strong enough immune responses to provide protection from infection, both RTS,S and R21 require at least three immunizations and subsequent boosters are critical to maintain efficacy^11^. Clinical efficacy is also not optimal. Phase 3 trials of RTS,S plus the adjuvant AS01 (RTS,S/AS01 _E_) in young children demonstrated that its efficacy against clinical malaria was ∼30->80%, depending on the age group and whether the trial occurred in low, moderate, or high transmission areas, but this efficacy declined over time^12,13^. Phase 3 trials of R21 adjuvanted with Matrix-M (R21/Matrix-M) indicated that its efficacy at preventing clinical malaria was somewhat higher (65-75%)^14^. However, differences between the design and location of the trials make it difficult to definitively demonstrate that R21/Matrix-M provides superior protection compared to RTS,S/AS01_E_ ^11^. Although the approval of these vaccines is a remarkable achievement in controlling malaria infection, the limitations of these vaccines underscore the critical need for the development of more potent vaccines capable of inducing stronger and more durable protective responses in a larger fraction of the at-risk population, consistent with the goal of the WHO to develop a vaccine that provides at least 75% protection against clinical malaria by 2030^15^.

CSP is an attractive vaccine target because of its abundance on the surface of sporozoites, its critical role in mediating sporozoite infectivity, and the fact that it contains highly conserved domains, unlike other, more variable, malaria surface antigens^16–19^. CSP consists of an immunodominant central repeat (CR) region which contains multiple (usually >35) copies of a short tetrapeptide (NANP) major repeat sequence interspersed with four copies of a minor repeat sequence (NVDP)^17^. The CR region is flanked by a flexible N-terminal domain, which contains a heparan sulfate binding site for hepatocyte attachment, and a structured C-terminal domain, which contains a thrombospondin-like type I repeat. RTS,S and R21 both contain a truncated form of CSP, consisting of 19 repeats of the NANP major repeat tetrapeptide plus the C-terminal domain, fused to the Hepatitis B virus surface antigen (HBsAg). When expressed, this fusion protein assembles into a virus-like particulate antigen. RTS,S is composed of a mixture of unfused HBsAg (∼80%) and the CSP-HBsAg fusion protein (∼20%)^20^, whereas R21 is only comprised of the CSP-HBsAg fusion protein, and therefore displays *Pf*CSP epitopes at higher valency^21^. Both RTS,S and R21 largely induce antibodies against the major repeat tetrapeptide because this sequence is immunodominant^22^. Although antibodies that bind to the NANP major repeat sequence can protect from malaria infection, achieving protective immunity requires an exceptionally high concentration of these antibodies^19,23,24^. The isolation of rare human monoclonal antibodies (mAbs) with potent ability to prevent malaria infection have identified new vulnerable epitopes within the CSP. These vulnerable epitopes lie at the CSP cleavage site, in the junctional region of CSP, between the N-terminal domain and the CR region, and within the major/minor tetrapeptide repeat region at the N-terminus of the CR region^25,26^. The most potent of these antibodies is L9 mAb^27^, which is notable for its ability to provide potent sterilizing immunity in passive immunization studies. This efficacy has been demonstrated in a mouse malaria infection model^27^, as well in experimental and natural infections in humans. Passive administration of L9 mAb was 88% effective in preventing malaria against controlled *Pf* infection^28^ and 70% effective at preventing natural *Pf* infection over a 6 month period in a field study in Mali^29^. These data suggest that targeting the vulnerable L9 epitope could be a promising strategy for producing next-generation vaccines with improved efficacy against malaria.

Previously, we engineered a VLP-based vaccine in which a synthetic peptide representing the 15 amino acid L9 epitope was linked at high valency to the surface of Qβ VLPs using a bifunctional chemical crosslinker. Immunization with Qβ L9 VLPs, in combination with Advax-3 adjuvant, elicited high titer antibody responses that resulted in sterilizing immunity in 60% of vaccinated mice in a malaria challenge model^30^. Qβ L9 VLPs display the L9 epitope as an unconstrained, linear epitope, raising the possibility that displaying L9 in a structural context that better mimics its native conformation could elicit more potent protective antibodies against *Pf*. Although the complete structure of *Pf*CSP remains unsolved, Martin and colleagues used cryo-EM to solve the structure of the L9 mAb in complex with recombinant *Pf*CSP^31^. This structure revealed that three L9 mAb-derived Fabs could bind to a 26 amino acid sequence encompassing three repeats of the motif NPNVDPNA. Each L9 Fab primarily contacted a core NPNV L9 mAb epitope, which adopts a type 1 beta-turn. The larger 26 amino acid epitope forms an extended loop structure when bound by three L9 Fabs. Given this structure, we asked whether display of the L9 epitope in a constrained loop on the surface of an immunogenic vaccine platform could potentially enhance the protective potency of elicited antibodies.

In this manuscript, we evaluate the immunogenicity and efficacy of recombinant MS2 VLPs displaying different lengths of the L9 mAb epitope in an exposed surface loop on the VLP surface. We show that these vaccines elicit potent and protective anti-CSP antibody responses, especially in combination with the adjuvant Cquim-MA (ViroVax), a dual TLR7/8 agonist-based adjuvant designed to traffic to draining secondary lymphoid tissue without systemic exposure. A combination vaccine, consisting of Qβ L9 and MS2 L9 VLPs adjuvanted with Cquim-MA was especially protective. Importantly, VLP-based vaccines targeting the L9 epitope were more effective than RTS,S/AS01_E_ at protecting mice from malaria.

## RESULTS

### Construction and assembly of MS2 VLPs displaying L9 peptides

Previously, we engineered the coat protein of bacteriophage MS2 so that it can tolerate insertion of heterologous peptide sequences in an exposed loop on its surface. This display site, which is referred to as the AB loop, consists of a beta-hairpin structure that protrudes from the surface of the virus particle (Fig. **1a**). Given that the core NPNV L9 epitope adopts a type 1 beta-turn^31^, we asked whether displaying this epitope in a more relevant structural context could elicit more potent antibody responses. Nucleotide sequences encoding peptides representing a minimal 8-amino acid L9 epitope centered around the core NPNV sequence (CSP amino acids 105-112), the consensus 15-amino acid epitope (amino acids 105-119, which contains two NPNV sequences), and an extended 27 amino acid epitope (amino acids 105-131) (Fig. **1b**) were inserted into the downstream AB-loop of the MS2 coat protein single-chain dimer expression vector. Expression plasmids were then used to express recombinant MS2 L9 VLPs in *E. coli*. Because each VLP consists of 90 copies of the single-chain dimer, each recombinant MS2 VLP displays exactly 90 copies of the L9 peptide per VLP. Successful insertion of the L9 peptides was confirmed by SDS-PAGE analysis. Coat proteins from recombinant MS2 L9 VLPs displayed a higher molecular weight compared to wild-type MS2 coat protein (Fig. **1c**). To confirm that the insertion of different L9 peptides into the MS2 coat protein was compatible with VLP assembly, transmission electron microscopy (TEM) was used to visualize the structures of the three different MS2 L9 VLPs (Fig. **1d**). These results showed that all three of the MS2 L9 constructs assembled into particles with diameters of approximately 30 nm. The morphology of MS2 VLPs displaying the longer 15- and 27-amino acid L9 sequences was somewhat less regular than VLPs displaying the shorter 8 amino acid peptide. Dynamic Light Scattering analysis of the recombinant MS2 L9 VLPs revealed that longer L9 insertions were associated with modest increases in the mean VLP diameter (Supplementary Figure 1). Of note, the 27aa L9 sequence is the largest peptide that we have successfully displayed on MS2 VLPs.

**Fig. 1.**
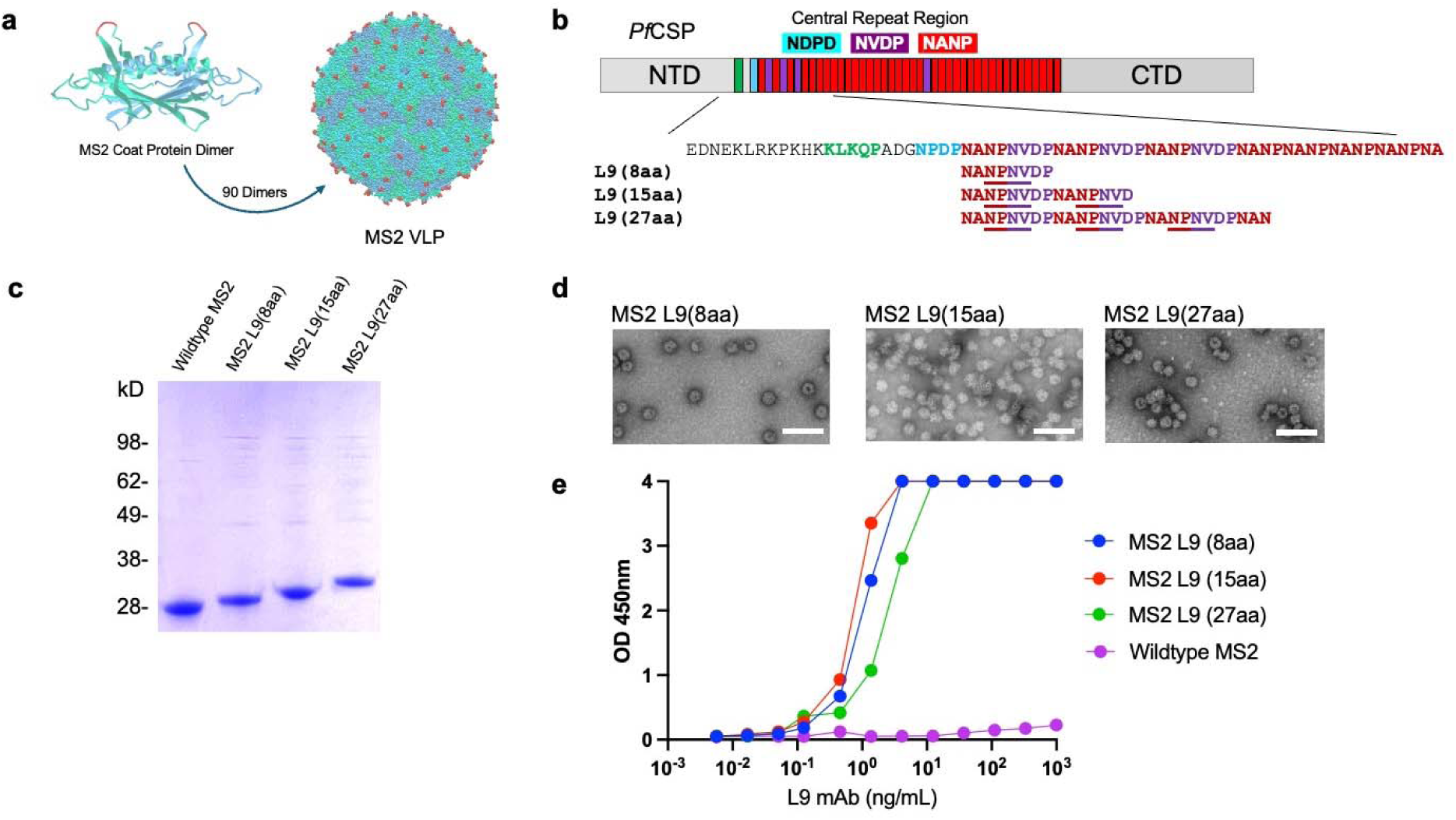
Characterization of MS2 L9 VLPs. **a** Structure of the MS2 coat protein dimer (left) and the MS2 VLP (right). 90 coat protein dimers self-assemble into a VLP. The location of the AB-loop is highlighted in red on both structures. Images of the MS2 coat protein dimer and the MS2 VLP were generated by the authors using online tools available through the RCSB protein data bank. **b** *Pf*CSP domains and the sequence of the junctional and minor repeat regions. The sequence of the three inserted L9 sequences and the location of the core NPNV epitopes (underlined) are shown. **c** SDS-PAGE analysis of wildtype MS2 VLPs and MS2 L9 VLPs displaying the 8, 15, and 27 amino acid L9 epitopes. The unmodified gel is shown in Supplementary Figure 2. **d** Transmission electron microscopy (TEM) images of recombinant MS2 L9 VLPs. The scale bar (in white) represents 100 nm. **e** Binding of L9 mAb to MS2 L9 VLPs or wildtype MS2 VLPs, as measured by ELISA.

The antigenicity of MS2 L9 VLPs was assessed by measuring the binding of L9 mAb to the MS2 L9 VLPs by enzyme-linked immunosorbent assay (ELISA). All three MS2 L9 VLPs were strongly recognized by the L9 mAb (Fig. **1e**), whereas wild-type MS2 VLPs were not. These findings indicate that displaying L9 epitopes in a constrained loop structure on MS2 VLPs does not affect L9 mAb binding.

### MS2 L9 VLPs elicit strong anti-CSP antibody responses

Next, we compared the immunogenicity of the three MS2 L9 VLPs constructs in mice. Mice were initially immunized twice, at weeks 0 and 3, and sera were collected one week after the second immunization to measure antibody levels against full-length recombinant CSP by ELISA. All three MS2 L9 VLPs elicited strong anti-CSP IgG responses (Fig. **2a**). MS2 L9(15aa) VLPs elicited the highest titer anti-CSP antibody responses. Based on their improved immunogenicity compared to the VLPs displaying the L9(8aa) epitope, MS2 L9 VLPs displaying the longer (15 and 27aa) epitopes were selected for further investigation. In each of these groups, mice received a third immunization to determine whether an additional boost would further enhance the immune response. This third immunization resulted in a substantial increase in anti-CSP antibody levels in mice that received the MS2 L9(27aa) VLPs, and a modest increase in anti-CSP antibody levels in mice immunized with MS2 L9(15aa) VLPs (Fig. **2b**). After three immunizations, the antibody responses induced by the MS2 L9(27aa) and MS2 L9(15aa) VLPs were nearly identical (Fig. **2b**).

**Fig. 2.**
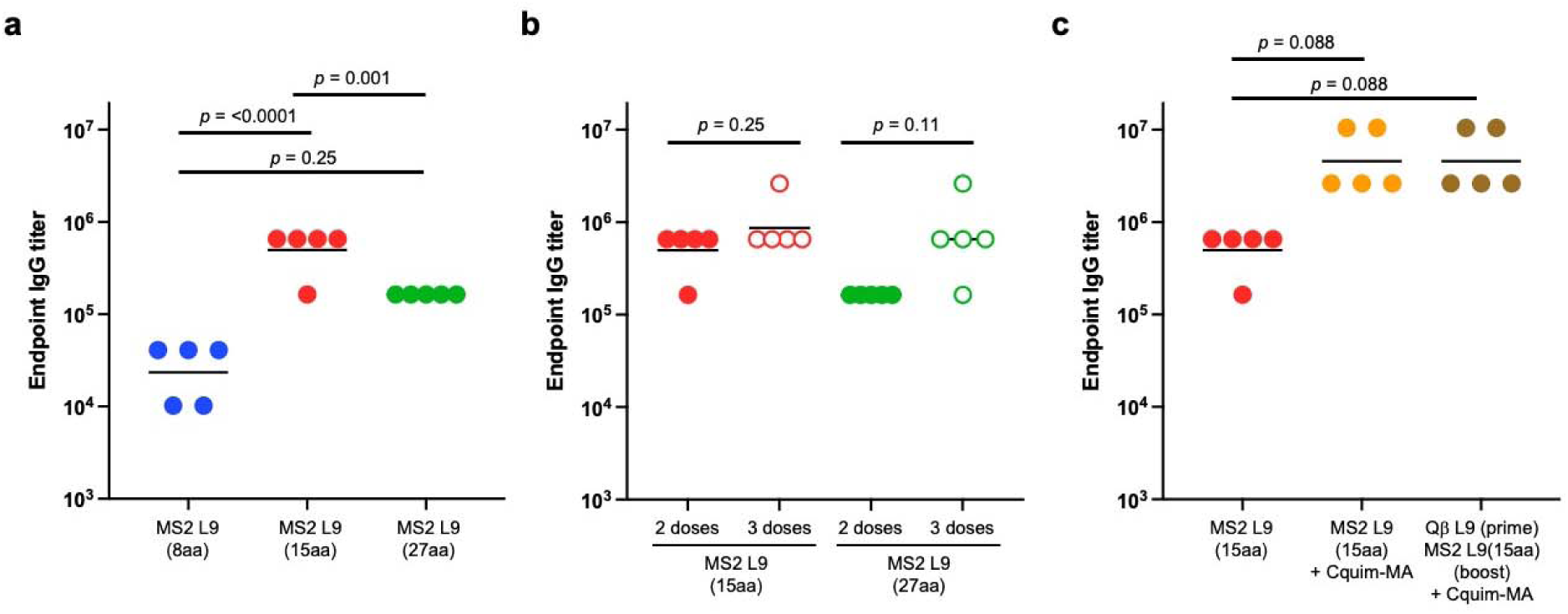
MS2 L9 VLPs elicit strong anti-CSP antibody responses. **a** Comparison of the immunogenicity of MS2 L9 VLPs displaying 8, 15, and 27 amino acid L9 epitopes. Groups of Balb/c mice (n=5) were immunized intramuscularly with 5 µg of VLPs at weeks 0 and 3. Anti-CSP IgG end-point dilution titers were calculated by ELISA using sera collected one week after the second immunization. Results show antibody titers in individual mice, lines represent the geometric mean titer from each group. Statistical comparisons were performed using a one-way ANOVA with post-hoc analysis. **b** Mice immunized with MS2 L9(15aa) and MS2 L9(27aa) received a third immunization, given 7 weeks after the prime. Sera was obtained one week following the third immunization. Statistical comparisons between titers following the second dose and the third dose were performed using an unpaired t-test. **c** Anti-CSP antibody titers in mice immunized twice with unadjuvanted MS2 L9(15aa) VLPs compared to mice that received two doses MS2 L9(15aa) plus Cquim-MA adjuvant and mice that received a Qβ L9 prime (plus Cquim-MA) followed by an MS2 L9(15aa) boost (also with Cquim-MA). Sera was obtained one week following the second immunization. Statistical comparisons were performed using a one-way ANOVA with post-hoc analysis (Tukey’s test).

We have previously shown that the immunogenicity and protective efficacy of Qβ VLP-based vaccines targeting CSP epitopes can be enhanced by co-administration of specific adjuvants. To evaluate if adjuvants could similarly enhance antibody responses to MS2-based vaccines, we measured the anti-CSP antibody responses generated by MS2 L9(15aa) VLPs formulated with Cquim-MA, a dual TLR7/8 agonist adjuvant, that we previously have shown can enhance antibody responses to Qβ VLP-based vaccines^32^. Two immunizations with MS2 L9(15aa) VLPs plus Cquim-MA induced markedly stronger anti-CSP antibody responses compared with unadjuvanted MS2 L9(15aa) VLPs (Fig. **2c**).

Heterologous prime-boost regimens can be an effective method for focusing antibody responses against specific epitope targets and minimizing antibody responses against the vaccine platform^33^. To determine if this approach could also be used to boost anti-CSP antibody responses, mice were primed with Qβ L9 VLPs mixed with Cquim-MA and then boosted with MS2 L9(15aa) VLPs, also formulated with Cquim-MA. This heterologous prime-boost approach resulted in similar anti-CSP antibody responses compared to those induced by MS2 L9(15aa) VLPs with Cquim-MA (Fig. **2c**). Taken together, these data demonstrate that MS2 L9 VLPs can elicit high titer anti-CSP antibody responses with or without heterologous boosting, especially in combination with Cquim-MA.

### MS2 L9 VLPs elicit durable antibody responses

One advantage of VLP-based vaccines is that they reliably elicit durable antibody responses^34^, likely through the efficient induction of long-lived plasma cells^35^. To assess the duration of the anti-CSP antibody response elicited by MS2 L9 VLPs, mice received three immunizations of MS2 L9(15aa) or MS2 L9(27aa) VLPs and antibody responses were measured for nearly a year following the prime. Both vaccines elicited long-lived antibodies, with little to no reduction in titers over 40-50 weeks following the initial prime (Fig. **3a**), mirroring what we previously observed following vaccination with Qβ L9 VLPs^31^. Similarly, we compared the durability of antibody responses in mice that received two doses of Cquim-MA adjuvanted vaccines (Fig. **3b**). Mice received either a homologous prime-boost with MS2 L9(15aa) VLPs or a heterologous prime-boost with Qβ L9 followed by MS2 L9(15aa) VLPs. Both vaccine regimens elicited similar peak antibody titers following the second immunization. Antibody levels slightly declined from the peak titer to the final timepoint, 38 weeks following the prime. Use of Cquim-MA with MS2 L9(15aa) VLPs slightly increased antibody titers compared to unadjuvanted vaccine at the late (38 week) timepoint, but this difference was not statistically significant.

**Fig. 3.**
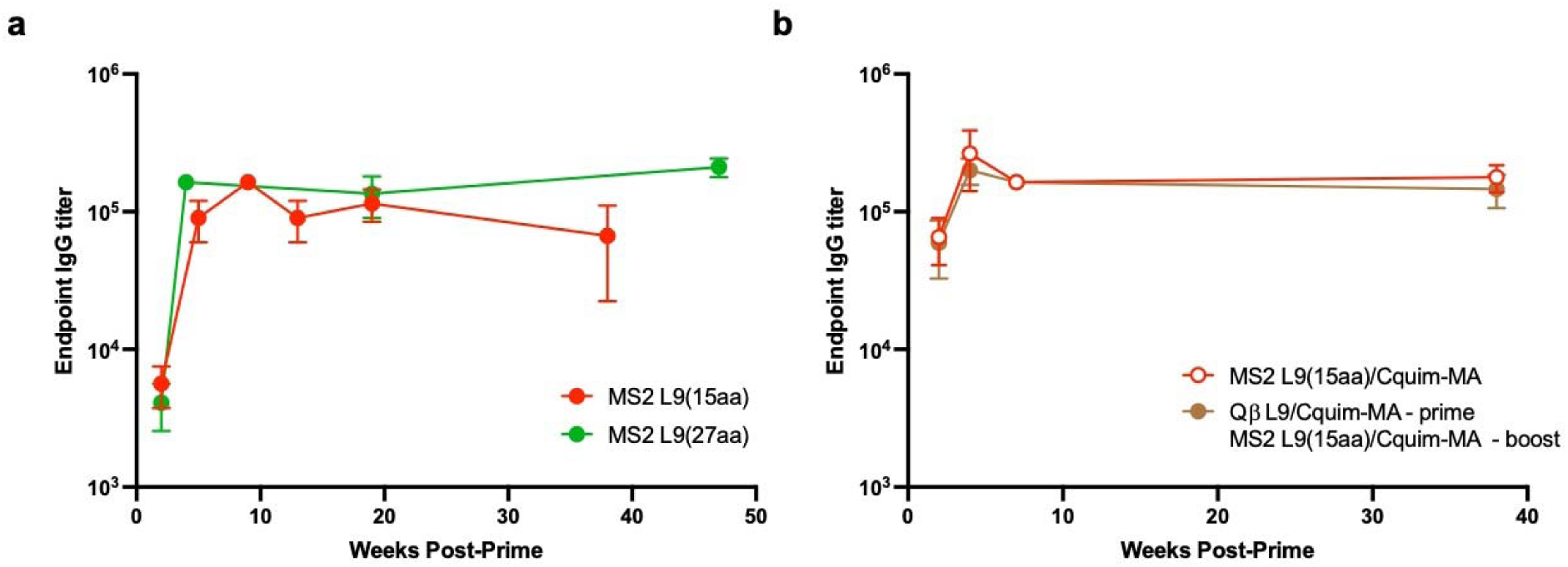
Longevity of antibody responses. **a** Groups of Balb/c mice (n=5) were immunized intramuscularly with 5 µg of VLPs at weeks 0, 3, and 7 and were measured following each immunization and at 38-47 weeks post-prime. Anti-CSP IgG titers were calculated by end-point dilution ELISA. Geometric means plus SEM are shown at each timepoint. **b** Mice that received vaccine plus Cquim-MA adjuvant received two immunizations, at weeks 0 and 3.

### Immunization with MS2 L9(15aa) VLPs plus Cquim-MA adjuvant protects mice from malaria challenge

Next, we evaluated whether MS2 L9 VLPs could protect mice from malaria challenge. Vaccine efficacy was assessed in a model in which vaccinated mice were challenged with mosquitoes carrying transgenic *Plasmodium berghei* (*Pb*) sporozoites expressing full-length *Pf*CSP and luciferase (*Pb*-*Pf*CSP-Luc)^37^. In this model, parasite liver loads can be quantified by measuring luciferase signal in the liver. In addition, the ability of vaccination to mediate sterilizing immunity can be determined by monitoring the development of blood-stage infection (parasitemia). In our initial challenge experiment (shown schematically in Fig. **4a**), we evaluated whether immunization with MS2 L9(15aa) VLPs could protect mice from malaria infection. Mice received three immunizations with MS2 L9(15aa) VLPs, with or without Cquim-MA. To benchmark vaccine efficacy, groups of mice were also immunized three times with Qβ L9 VLPs plus Cquim-MA or with the WHO-approved vaccine RTS,S/AS01_E_ (using a 5 µg dose). As negative controls, groups of mice were immunized with wild-type MS2 VLPs (not displaying the L9 epitope), wild-type Qβ VLPs, or Cquim-MA adjuvant alone, or were unimmunized (naïve). Following immunization, anti-CSP antibody concentrations from individual mice were quantified. As shown in Fig. **4b**, RTS,S/AS01_E_, Qβ L9/Cquim-MA, MS2 L9(15aa), and MS2 L9(15aa)/Cquim-MA induced strong anti-CSP antibody responses. MS2 L9(15aa)/Cquim-MA elicited the highest anti-CSP antibody concentrations, with mean levels ∼6-fold higher than unadjuvanted MS2 L9(15aa), ∼3-fold higher than Qβ L9/Cquim-MA, and ∼2.2-fold higher than RTS,S/AS01_E_. None of the control groups elicited anti-CSP antibodies (not shown).

**Fig. 4.**
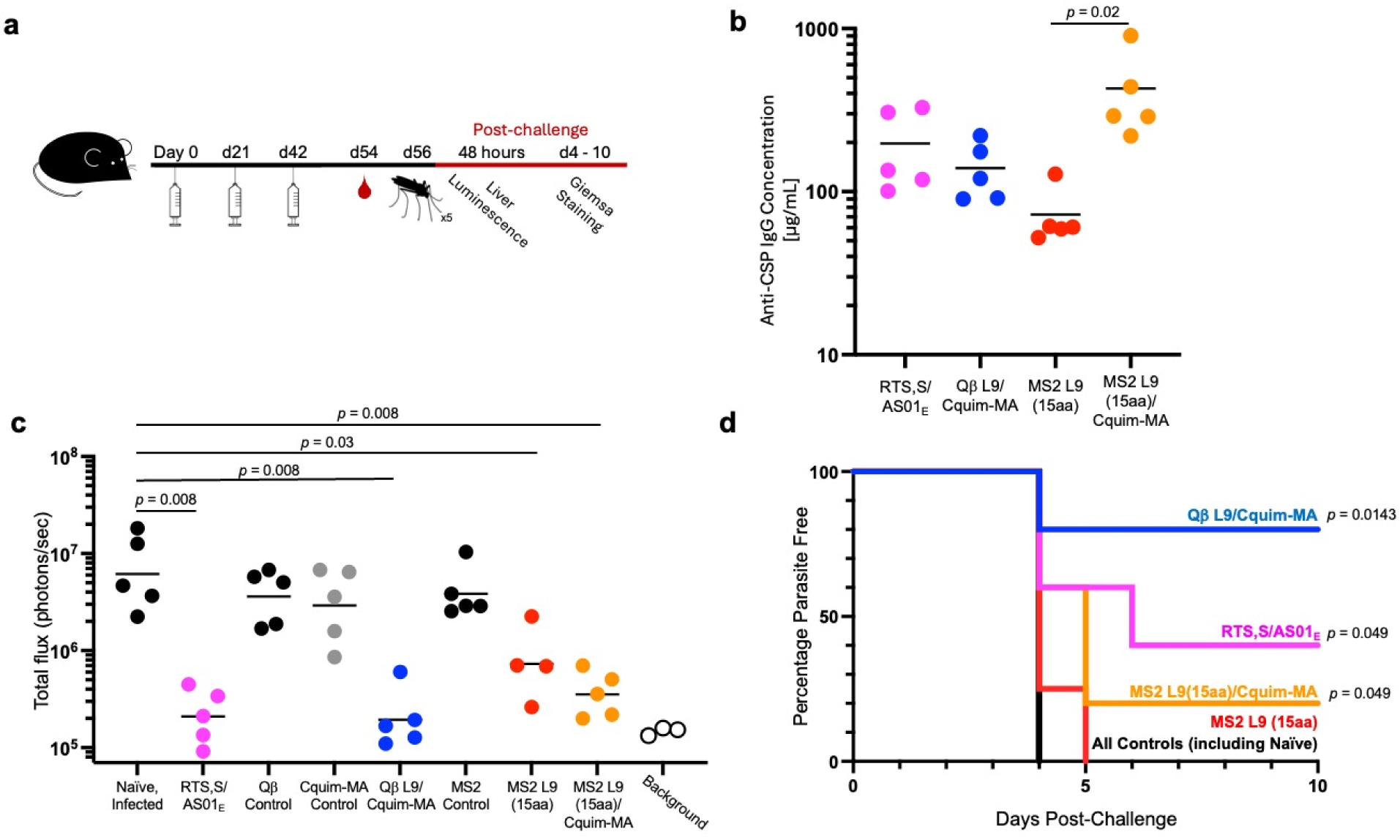
Both Qβ L9 and MS2 L9(15aa) VLPs plus Cquim-MA elicit strong antibody responses and protect against malaria infection. **a** Experimental timeline. C57BL/6 mice were immunized three times at weeks 0, 3, and 6, followed by a challenge with five *Pb-PfCSP-Luc*-infected mosquitoes at week 8. Liver luminescence was measured 42 hours after the mosquito challenge, and blood smears were taken starting on day 3 post-infection. **b** Anti-CSP antibody levels in serum obtained following the third vaccination, prior to challenge. Horizontal lines represent the mean anti-CSP antibody concentration for each group. Groups are compared by 2-tailed unpaired *t* test. **c** Parasite liver burden measured via luminescence. Horizontal lines indicate geometric mean luminescence for each group. Statistical comparisons were performed using a two-tailed Mann Whitney test. **d** Protection from blood-stage infection, as measured by percent of blood parasite-free mice post-challenge. A log-rank test (controlling for multiple comparisons) was used to statistically compare protected groups to naïve mice.

Mice were then infected by exposure to five *Pb*-*Pf*CSP-Luc infected mosquitoes. As an initial determination of vaccine efficacy, liver-stage parasite burden was quantified by measuring liver luciferase activity using intravital imaging 42 hours post-infection (Fig. **4c**). As has been previously shown in this model^36^, immunization with RTS,S/AS01_E_ dramatically reduced parasite liver burden, by ∼97% relative to naïve controls. Immunization with Qβ L9/Cquim-MA also reduced parasite liver burden by ∼97%. Groups immunized with MS2 L9(15aa) and MS2 L9(15aa)/Cquim-MA also displayed statistically significant reductions in parasite liver loads (by 88% and 94%, respectively). None of the negative control groups were protected from infection.

To evaluate the possibility of sterilizing protection, daily blood smears were taken from the challenged mice starting at day 3 post-infection. Blood was assessed for parasitemia using Giemsa staining (Fig. **4d**). While all control groups developed blood-stage parasitemia by day 4, 40% of the RTS,S/AS01_E_-vaccinated group (2 out of 5 mice) were protected, which is similar to what has been previously reported in this model^36^ and mirrors the protection conferred by RTS,S/AS01_E_ in humans. 80% of the mice immunized with Qβ L9/Cquim-MA were completely protected from infection, which is an improvement compared to previous studies using Qβ L9 adjuvanted with Advax-3^30^, suggesting that Cquim-MA may be a more effective adjuvant in combination with this vaccine. In this challenge study, immunization with Qβ L9/Cquim-MA provided stronger protection compared to RTS,S/AS01_E_ (80% compared to 40%), but this difference was not statistically significant (*p*=0.239). Although MS2 L9(15aa)/Cquim-MA elicited the highest anti-CSP antibody concentrations, only 20% of mice immunized with this vaccine remained free from blood-stage parasitemia.

### A combination L9 VLP vaccine reduces liver parasite burden and enhances sterilizing immunity in *Plasmodium*-challenged mice

We performed a second, independent challenge experiment with the following goals: (1) to replicate our initial results showing that vaccination with Qβ L9/Cquim-MA provides stronger protection than RTS,S/AS01_E_, (2) to evaluate the protective efficacy of MS2 L9 VLPs displaying the longer 27aa L9 peptide, and (3) to assess protection of a combination of the MS2 L9(15aa) and Qβ L9 vaccines. This experiment followed the same design as our initial study; mice were immunized with RTS,S/AS01_E_, Qβ L9/Cquim-MA, MS2 L9(27aa)/Cquim-MA, or a mixture of Qβ L9 and MS2 L9(15aa)/Cquim-MA. Negative controls included a group of mice that received wildtype Qβ VLPs and a group of unimmunized (naïve) mice. As we showed previously, Qβ and MS2-based L9 vaccines elicited strong anti-CSP antibody responses, with mean levels >100 µg/mL, similar to the antibody levels induced by RTS,S/AS01_E_ (Fig. **5a**). Immunization with Qβ L9/Cquim-MA or MS2 L9(27aa)/Cquim-MA resulted in similar reductions in parasite liver loads as RTS,S/AS01_E_ (∼91%), but mice that received the combination vaccine plus Cquim-MA showed the greatest reduction in parasite burden (∼96%), a reduction that was statistically distinct from RTS,S/AS01_E_ (*p*=0.03) (Fig. **5b**). Similar to our initial challenge experiment (Fig. **4**), vaccination with Qβ L9/Cquim-MA resulted in sterilizing immunity in a higher percentage of mice than we observed in the group immunized with RTS,S/AS01_E_ (Fig. **5c**), although this difference was not statistically significant (*p*=0.25). The percentage of mice that were protected from infection were lower in this experiment (43%) compared to the previous challenge study (80%). This may be explained by the higher mean parasite liver burden across all groups in the second challenge experiment (for example, it was 20% higher the naïve group) or it could reflect variability of parasite burden in this challenge model. Interestingly, immunization with the combination vaccine (Qβ and MS2 L9 VLPs with Cquim-MA) provided the strongest protection, with a higher percentage of mice exhibiting sterilizing immunity than in mice immunized with either RTS,S/AS01_E_ or Qβ L9/Cquim-MA. Taken together, these results indicate that a combination vaccine presenting the L9 epitope in multiple conformations, as an unstructured peptide on Qβ VLPs and in a structured β-hairpin on MS2 VLPs, was effective at eliciting sterilizing immunity against *Plasmodium* infection.

**Fig. 5.**
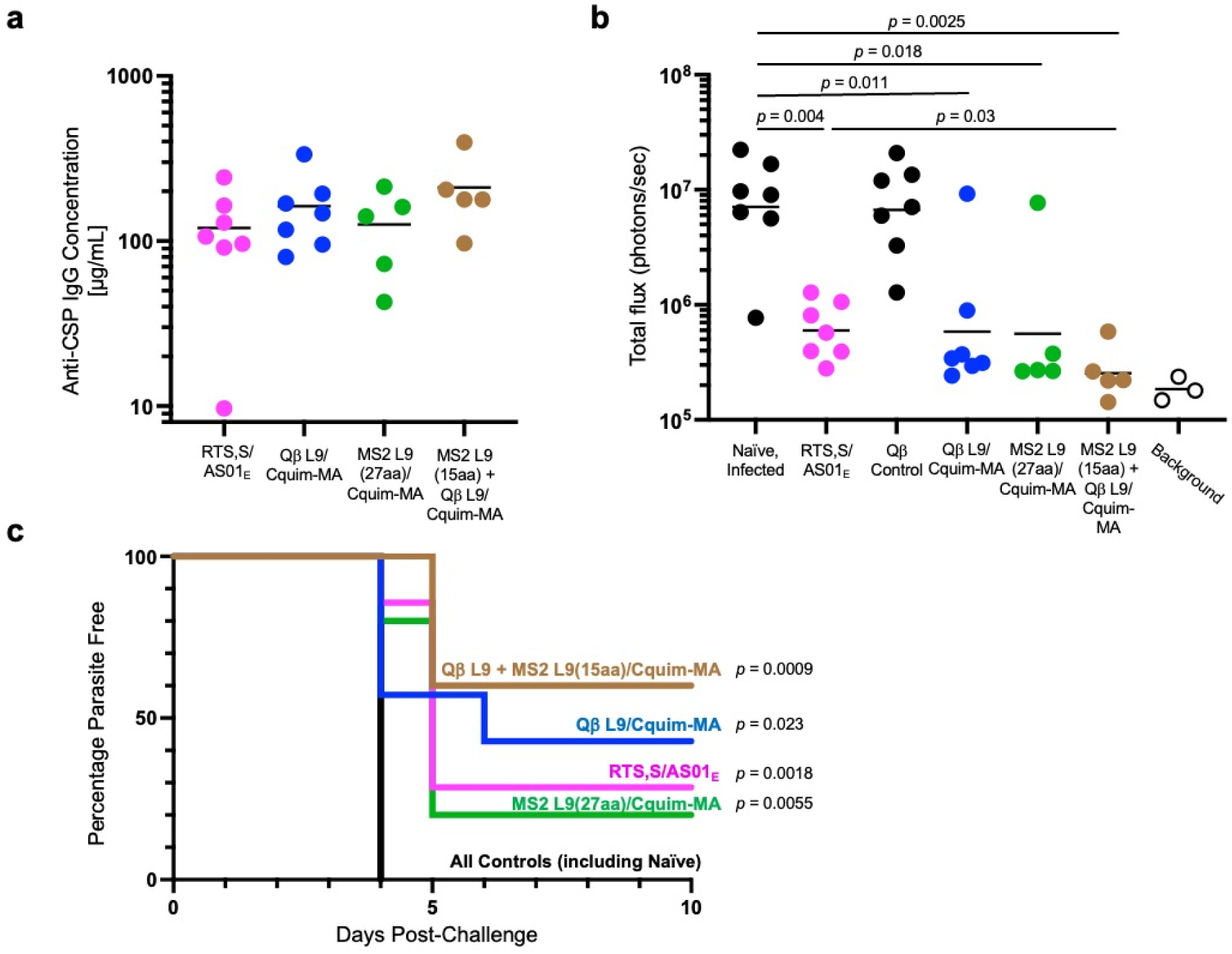
A combination vaccine targeting the L9 epitope provides the strongest protection from malaria infection. **a** Anti-CSP antibody levels in serum obtained following the third vaccination. Horizontal lines represent the mean anti-CSP antibody concentration for each group. No significant differences were observed between groups (p > 0.05, unpaired t-test). **b** Parasite liver burden measured via luminescence. Horizontal lines indicate geometric mean luminescence for each group. Statistical comparisons were performed using a two-tailed Mann Whitney test. **c** Protection from blood-stage infection, as measured by percent of blood parasite-free mice post-challenge. A log-rank test (controlling for multiple comparisons) was used to statistically compare protected groups to naïve mice.

## DISCUSSION

The identification of potent mAbs against *Pf*CSP have helped guide efforts to develop next-generation vaccines with improved efficacy against malaria infection. These mAbs include MAM01, which targets the major NANP repeats in the central repeat^37^, CIS43, which preferentially targets a unique sequence (DPNA) in the junctional region^38^, and L9, which preferentially binds to NPNV sequences in the minor repeat region^27^. Of these mAbs, L9 has the highest potency in preventing malaria infection in mouse models, approximately 3-fold better than CIS43^27^. Correspondingly, we have shown that a Qβ VLP-based vaccine displaying the L9 epitope had increased efficacy compared to a similar vaccine targeting the CIS43 epitope^30^. In this study, we evaluated whether the structural context of the L9 epitope and whether a novel adjuvant, Cquim-MA, could enhance vaccine efficacy.

VLP-based vaccines that present the L9 epitope as an unconstrained, linear epitope are capable of eliciting high-titer protective antibodies against malaria^30^. However, when bound to the L9 mAb, the core NVDP L9 epitope adopts a type 1 beta-turn^31^, suggesting that displaying the L9 epitope in a similar structural context could elicit more potently protective antibodies. We produced recombinant MS2 bacteriophage VLPs in which L9 epitopes of different lengths were inserted into a naturally occurring beta-hairpin loop that is prominently displayed on the surface of the VLP. MS2 L9 VLPs, particularly those displaying the longer 15- and 27-amino acid L9 peptides, elicited robust antibody responses against *Pf*CSP. When used in conjunction with Cquim-MA adjuvant, MS2 L9(15aa) VLPs, in particular, elicited remarkably strong anti-CSP antibody levels, approximately 4-fold higher than Qβ L9/Cquim-MA and 2-fold higher than RTS,S/AS01_E_. MS2 L9 VLPs were also able to induce durable immunity; there was only a minimal decline in antibody titers in mice for nearly a year following immunization. This is consistent with previous studies showing the longevity of antibody responses to VLP-based vaccines, which are known to elicit long-lived plasma cells^34,35^.

We previously showed that the adjuvant Advax-3, which is a co-formulation of CpG55.2 oligonucleotide with aluminum hydroxide, could increase anti-CSP antibody levels induced by Qβ CIS43 VLPs by approximately 8-fold^39^. However, over time, antibody concentrations in mice immunized Qβ CIS43 VLPs plus Advax-3 declined to similar levels as mice immunized with unadjuvanted vaccine. Here, we showed that the novel dual TLR7/8 agonist Cquim-MA could strongly boost antibody responses to both Qβ L9 and MS2 L9 VLPs. These antibody responses were also durable; we observed only a slight reduction in antibody titers over a period of 9 months following the primary immunization. The ability to elicit durable antibody responses is likely to be particularly important for providing protection from malaria infection, especially given the brief window between infection and hepatocyte invasion in which sporozoites exist extracellularly^40^.

To assess vaccine efficacy, we evaluated vaccine candidates in a well-established mouse malaria challenge model that was designed to evaluate the efficacy of *Pf*CSP-targeted vaccines^41^. Importantly, we were able to benchmark VLP-based vaccines against the WHO-approved vaccine, RTS,S/AS01_E_. The Zavala laboratory and PATH had previously developed a standardized model for comparing the potency of next-generation vaccines to RTS,S/AS01_E_ ^36^, and we adopted this model in our study. Control mice received three immunizations of RTS,S/AS01 at a dose (5 µg) that had been shown to maximize anti-CSP antibody responses^36^. While MS2 L9 VLPs/Cquim-MA elicited very strong antibody responses, these vaccines were less effective than Qβ L9/Cquim-MA or RTS,S/AS01_E_ at reducing parasite liver burden and conferring sterilizing immunity. Qβ L9 VLPs formulated with Cquim-MA elicited similar antibody levels as RTS,S/AS01_E_, but consistently provided stronger protection. Whereas vaccination with RTS,S/AS01_E_ conferred sterilizing immunity in 28-40% of mice, which is slightly lower efficacy than was previously reported^36^, immunization with Qβ L9/Cquim-MA protected 43-80% of vaccinated mice. Importantly, both vaccines provided protection and the differences in efficacy between the two vaccines were not statistically significant. Nevertheless, these data provide strong support for continued focus on the L9 epitope as a particularly vulnerable target for protective antibodies.

Interestingly, a combination vaccine consisting of MS2 L9 and Qβ L9 VLPs demonstrated strong protection from infection. Although these protection data will need to be repeated to confirm this observation, the preliminary efficacy of this mixed vaccine highlights the potential of combining antigens presented in different conformational contexts—unstructured epitopes on Qβ VLPs and structured β-hairpins on MS2 VLPs. The flexible central repeat of CSP is intrinsically disordered, adopting multiple conformations^42,43^. Thus, the improved protection may stem from the ability of these vaccines to activate B cell subsets that recognize and respond to the L9 epitope in multiple forms, thus broadening the immune response and increasing the likelihood of robust protection. Additional investigation of the structure of the synthetic L9 peptide displayed on Qβ VLPs could potentially provide insight on the structural basis for these differences in functional antibody responses.

While the combination vaccine outperformed individual VLP platforms, sterilizing immunity was not observed in all vaccinated mice, emphasizing the challenges still faced in developing a fully protective malaria vaccine. However, the success of this approach suggests that further refinement of the vaccine, potentially by optimizing epitope presentation or exploring additional adjuvants, could lead to even stronger and more consistent sterilizing protection. Taken together, these results highlight the potential of multi-platform VLP vaccines in providing effective, durable, and sterilizing immunity against malaria. The success of combining MS2 and Qβ VLPs to present the L9 epitope in multiple conformations represents a promising strategy for malaria vaccine development, although there are also complications associated with development and deployment of two-component vaccines. Further development of L9 epitope-based VLP vaccines, in combination with those that elicit antibody and/or T-cell responses against other liver-stage malarial antigens, could broaden and enhance the overall protective efficacy against malaria.

## METHODS

### Production and Characterization of Recombinant MS2 VLPs

The MS2 VLP expression plasmid pDSP62, which encodes the single-chain dimer version of the MS2 bacteriophage coat protein, was generated previously^44,45^. Sequences encoding the L9 epitope (representing CSP amino acids 105-112, 105-119, and 105-131) were generated by PCR and cloned into pDSP62 so that the L9 epitope was inserted into the surface-exposed AB-loop of the downstream copy of the MS2 bacteriophage coat protein. Cloned constructs were sequenced to confirm the presence of the sequence encoding the L9 epitope.

MS2 VLPs displaying L9 epitopes were expressed in *E. coli*. Expression plasmids were used to transform the C41 *E. coli* expression strain by electroporation. Transformed C41 cells were grown at 37°C using Luria Bertani broth containing 50 µg/mL kanamycin until the cells reached an OD600 of 0.6. Coat protein expression was induced using 0.4 mM isopropyl-β-D-1-thiogalactopyranoside and grown at 37°C overnight. Cell pellets were collected, re-suspended in lysis buffer [50 mM Tris-HCL, 100 mM NaCl, 10 mM ethylenediaminetetraacetic acid, pH 8.5, 0.45% deoxycholate] and incubated on ice for 30 min. Cells were lysed by 3-5 cycles of sonication at 20Hz and then cell lysates were clarified by centrifugation (15,000×*g*, 20 min, 4°C). Soluble MS2 L9 VLPs were purified by precipitation using 70% saturated (NH_4_)_2_SO_4_, followed by size exclusion chromatography (SEC) using a Sepharose CL-4B column. The column was pre-equilibrated with a purification buffer (40 mM Tris-HCl, 400 mM NaCl, 8.2 mM MgSO_4_, pH 7.4). Fractions containing VLPs were identified by agarose gel electrophoresis and/or SDS-PAGE. MS2 L9 VLPs were concentrated from SEC purified fractions by precipitation using 70% saturated (NH_4_)_2_SO_4_ followed by dialysis against PBS (pH 7.4). Endotoxin was removed from preparations by three rounds of Triton X-114 (Sigma-Aldrich) phase separation, using a published protocol^46^. The final concentration and purity of VLPs was determined via SDS-PAGE using a 10% NuPAGE gel (Invitrogen) using known concentrations of hen egg lysozyme (HEL) as standards. Protein concentrations were determined using ImageJ.

VLPs were visualized by transmission electron microscopy (TEM). VLPs were adsorbed on carbon-coated glow-discharged copper grids for 2 minutes and then were negatively stained with 2% uranyl acetate for 2 minutes. VLPs were visualized at a magnification of 100,000X.

The display of L9 epitopes on the surface of recombinant VLPs was confirmed by ELISA. Briefly, 250 ng of MS2 L9 VLPs were coated onto wells of an Immunulon 2 ELISA plate (Thermo Fisher Scientific) and probed with serial dilutions of the L9 monoclonal antibody (L9 mAb), generously provided by Robert Seder at the NIH Vaccine Research Center. L9 mAb binding was detected using a 1:4000 dilution of horseradish peroxidase (HRP)-conjugated goat anti-human IgG (Jackson Immunoresearch), followed by the addition of 3,3′,5,5′-tetramethylbenzidine (TMB) substrate (Thermo Fisher Scientific). The reaction was stopped by the addition of 1% HCl, and optical density was measured at 450 nm (OD450) using an AccuSkan plate reader (Fisher Scientific).

### Production of Qβ L9 VLPs

Qβ L9 VLPs were generated by chemically conjugating a synthetic peptide representing the CSP L9 epitope to preformed Qβ VLPs. A peptide representing the L9 epitope of CSP, modified to contain the C-terminal linker sequence *gly-gly-gly-cys* (NANPNVDPNANPNVD-*GGGC*), was synthesized by GenScript. L9 peptides were conjugated to exposed lysine residues on the surface of Qβ VLPs by using the heterobifunctional crosslinker succinimidyl 6-[(beta-maleimidopropionamido) hexanoate] (SMPH; Thermo Fisher Scientific). SMPH was incubated with Qβ VLPs at a molar ratio of 10:1 (SMPH:Qβ coat protein) for 2 hours at room temperature. Excess SMPH was removed using an Amicon Ultra-4 centrifugal unit with a 100 kDa cutoff (Millipore). L9 peptide was incubated with Qβ VLPs at a molar ratio of 10:1 (peptide:Qβ coat protein) and allowed to react overnight at 4°C.

### Dynamic Light Scattering (DLS)

Size measurements were conducted using a Malvern Zetasizer Nano ZS and VLPs at a concentration of 0.5–1 mg/mL. Each sample was tested in triplicate. Mean diameters for each sample were calculated using Zetasizer software from the volume-based distribution.

### Ethics Statement for Animal Studies

All animal research complied with the Institutional Animal Care and Use Committee at the University of New Mexico School of Medicine (approved protocol #: 22-201289-HSC) and at Johns Hopkins University (approved permit #: MO18H419).

### Mouse Immunization and Challenge Studies

To assess vaccine immunogenicity, groups of 4–5-week-old female Balb/c mice (typically n = 5 per group; Jackson Laboratory) were immunized intramuscularly in the ventral flank with 5 μg of VLPs or control VLPs in a volume no greater than 50 µL. Doses were calculated based on the total amount of protein in the preparation. Because there are less copies of the L9 peptide on MS2 VLPs (90 per VLP) compared to Q-beta VLPs (∼350 per VLP), the total dose of the L9 peptide was ∼4-fold lower in mice immunized with MS2 L9 VLPs. When adjuvant was used, vaccines were mixed with 5 µg of Cquim-MA, a dual TLR7/8 agonist generously provided by ViroVax, LLC. Mice were boosted with the same dose of vaccine and adjuvant three and seven weeks after the initial prime. Sera were collected by retroorbital bleed one week after each immunization and, in some cases, monthly thereafter. For sera collection, mice were sedated using isoflurane (2-4% with oxygen as a carrier for 3 minutes). At the conclusion of each study, mice were euthanized by terminal cardiac puncture. Animals were anesthesized by intraperitoneal administration of 0.3 mL of a mixture of ketamine (10 mg/mL) and xylazine (1 mg/mL) prior to the blood draw. Euthanasia was confirmed by cervical dislocation.

For challenge studies, groups of 7–8-week-old C57Bl/6 mice were used. Mice were immunized with 5 μg of VLPs with or without 2 µg Cquim-MA adjuvant. As negative controls, mice were immunized with wild-type (unmodified) VLPs, Cquim-MA adjuvant alone, or were unimmunized. As benchmark controls, mice received a 5 µg intramuscular dose of RTS,S with 10-fold diluted original AS01_E_ adjuvant dose, representing 2.5 µg of the TLR4 ligand 3-*O*-desacyl-4’-monophosphoryl lipid A (MPL), and 2.5 µg of the QS-21 saponin. Vaccine was administered on days 0, 21, and 42. Sera were collected following the final boost (at day 54) as described above and then mice were challenged two days later.

Mice were challenged using *Anopheles stephensi* mosquitoes infected with transgenic *P. berghei* sporozoites engineered to express luciferase and full-length *Pf*CSP in place of *P. berghei* CSP, denoted as Pb-PfCSP-luc^41^. Infected mosquitoes were generated by allowing the insects to blood-feed on Pb-PfCSP-Luc-infected mice. Prior to exposure, mice were anesthetized with 2% Avertin. Each mouse was then exposed to five infected mosquitoes for a 10-minute blood meal. After feeding, the number of mosquitoes with visible blood meals was recorded. Liver luminescence was measured 42 hours post-challenge. Mice were sedated using 2-4% isoflurane, and 100 μL of D-luciferin (30 mg/mL) was administered via intraperitoneal injection. Luminescence from the liver was subsequently captured using the IVIS Spectrum Imaging System (Perkin Elmer) while the mice were under sedation using isoflurane. Starting 4 days post-challenge, blood smears were collected daily and stained with Giemsa to monitor parasitemia.

### Quantification of Antibody Responses

Anti-CSP antibody responses were quantified by ELISA as reported previously^30,39^. To measure serum antibody levels against full-length CSP, an ELISA was performed using recombinant CSP, expressed in Pseudomonas fluorescens and kindly provided by Gabriel Gutierrez (Leidos, Inc.).^47^ Immulon 2 ELISA plates (Thermo Scientific) were coated with 250 ng of recombinant CSP in 50 μL of PBS per well overnight at 4°C or for 2 hours at room temperature (RT). Wells were blocked with 100 µL PBS-0.5% nonfat dry milk for 1 hour at RT or overnight at 4°C. Sera isolated from immunized mice were serially diluted in PBS-0.5% milk and applied to wells for 2 hours at RT or overnight at 4°C. Reactivity was measured by addition of HRP-labeled goat anti-mouse IgG, diluted 1:4000 in PBS-0.5% milk (Jackson Immunoresearch) for one hour at RT, and detected by addition of TMB substrate. Reactions were stopped using 1% HCl and optical density was measured at 450 nm (OD450) using an AccuSkan plate reader (Fisher Scientific). For some challenge experiments, CSP-specific antibody concentrations in mouse sera were quantified by linear regression analysis using a standard curve generated using known concentrations of the anti-CSP mouse monoclonal antibody 2A10, as described previously^30,39^.

### Statistical Analysis

All statistical analyses were conducted using GraphPad Prism v.9. Unpaired two-tailed t-tests were used for comparing antibody levels and Mann-Whitney tests were used for the mosquito challenge experiments. For percent inhibition calculations, liver luminescence values of individual vaccinated mice were divided by the mean of mice in negative control groups. Background luminescence levels were subtracted from all values.

### Data availability

All data generated or analysed during this study are included in this published article and its supplementary information files.

## ACKNOWLEDGEMENTS

This research was funded by the National Institutes of Health (R01 AI169739 to B.C.). The authors also thank a generous contribution to the UNM Foundation in honor of Jeffrey Michael Gorvetzian in support of biomedical research excellence at the University of New Mexico School of Medicine (to B.C.) and Bloomberg Philanthropies (to Y.F-G. and F.Z.). A.F. was supported by the UNM Academic Science Education and Research Training (ASERT) program (funded by K12 GM088021). S.D. (ViroVax LLC) gratefully acknowledges support from the NIAID (Adjuvant Development Contract HHSN272201800049C) which allowed the discovery and development of Cquim-MA. We also acknowledge facilities provided by the Autophagy, Inflammation, & Metabolism (AIM) Center of Biomedical Research Excellence (COBRE) cores, funded by NIH grant P20 GM121176, and the Insectary of the Johns Hopkins Malaria Research Institute for support. We also thank Emily Locke (PATH) for a critical reading of the manuscript, GSK for providing access to RTS,S/AS01_E_ and for comments on the manuscript, and Aidan Leyba, Dinesh Choudhury, and Pavan Muttil for assistance with DLS experiments.

## CONTRIBUTIONS

This work was conceived by B.C., Y.N., and F.Z. and A.F. Y.N. (all figures), A.F. (figures 4-5), B.R. (figure 1), and Y.F-G. (figures 4-5) conducted the experiments and data analysis, S.A.D. contributed critical reagents (Cquim-MA). The manuscript was drafted by Y.N. and B.C.; all authors edited the manuscript and approved its submission.

## Notes

### Competing Interest Statement

B. Chackerian has equity in Metaphore Biotechnologies. S. David is affiliated with ViroVax LLC, which holds proprietary rights over the Cquim-MA adjuvant. All other authors have no competing interests.

### Summary of Updates

The title has been shortened and the supplemental information has been expanded.

