## Supplementary Information for "Virus-like particle vaccines targeting a key epitope in circumsporozoite protein provide sterilizing immunity against malaria"

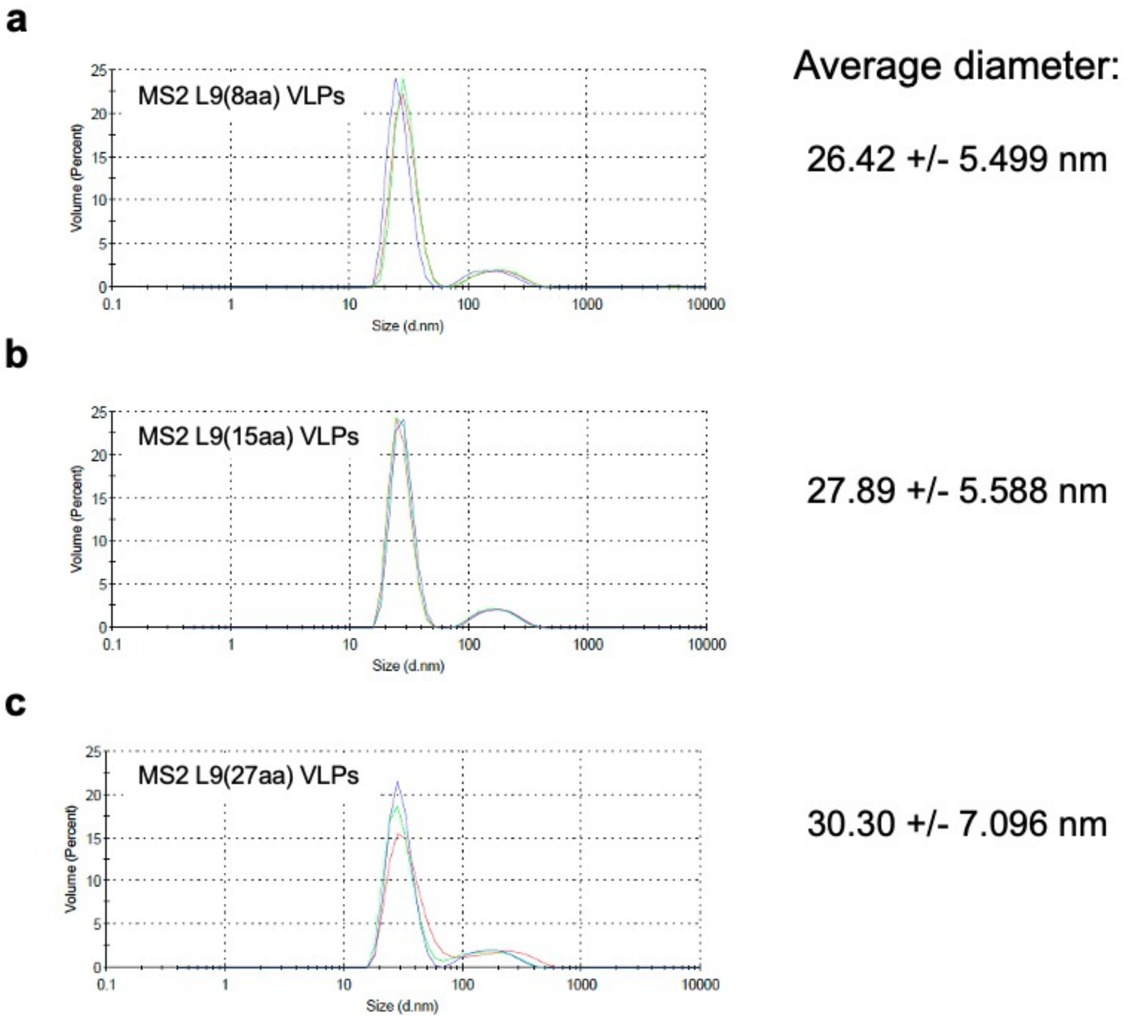


**Supplementary Figure 1. Dynamic Light Scattering (DLS) analysis of MS2 L9 VLPs.** DLS analysis of **a** MS2 L9(8aa), **b** MS L9(15aa), and **c** MS2 L9(27aa) VLPs. Experiments were performed in triplicate; each replicate is indicated on the graph using a different color line. Average VLP diameter +/- standard deviation (in nm) was calculated and is indicated to the right of the graphs.

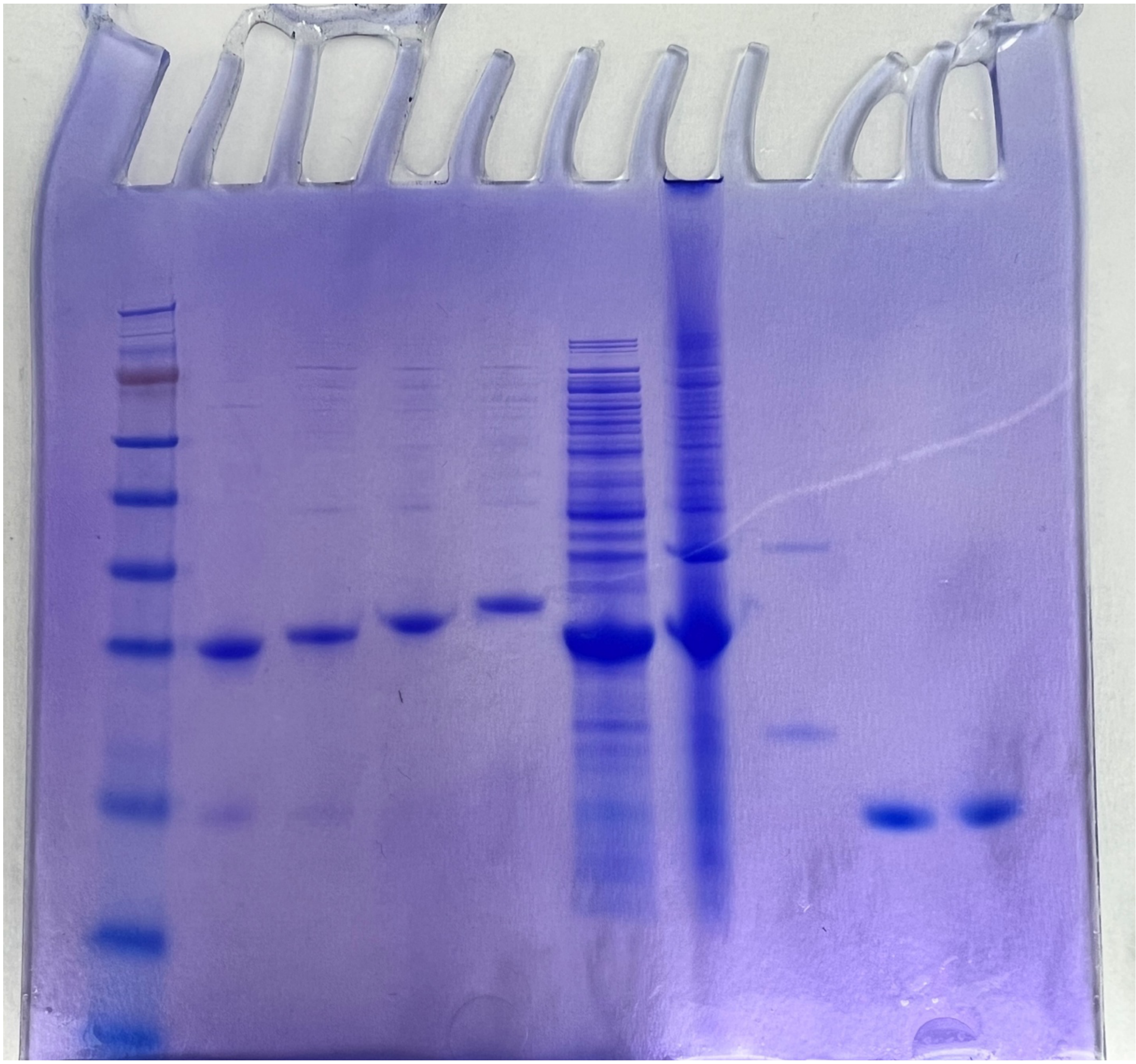

**Supplementary Figure 2.** Unmodified gel used as a source of the data shown in Figure 1c. Lane 1 (left) contains molecular weight markers (Invitrogen SeeBlue Plus2 Prestained Markers). Lane 2 contains wildtype MS2 single-chain dimer VLPs. Lane 3 contains MS2 L9(8aa) VLPs. Lane 4 contains MS2 L9(15aa) VLPs. Lane 5 contains MS2 L9(27aa) VLPs. Lanes 6-10 are samples from an unrelated experiment. Lanes 2-5 are shown in Figure 1c.
